# *In vivo* exchange of glucose and lactate between photoreceptors and the retinal pigment epithelium

**DOI:** 10.1101/2023.04.10.536306

**Authors:** Daniel T. Hass, Elizabeth Giering, John Y.S. Han, Celia M. Bisbach, Kriti Pandey, Brian M. Robbings, Thomas O. Mundinger, Nicholas D. Nolan, Stephen H. Tsang, Neal S. Peachey, Nancy J. Philp, James B. Hurley

## Abstract

Photoreceptors in the retina of a vertebrate’s eye are supported by a tissue adjacent to the retina, the retinal pigment epithelium (RPE). The RPE delivers glucose to the outer retina, consumes photoreceptor outer segments discs, and regenerates 11-cis-retinal. Here we address the question of whether photoreceptors also provide metabolic support to the RPE. We use complementary approaches and animal models to show that glucose is the primary fuel for the retina, that photoreceptors are the primary cell type in the retina to consume glucose, and that lactate derived from photoreceptor glucose consumption is transported to and catabolized by the RPE. These data rigorously support and extend the concept of a metabolic ecosystem between photoreceptors and RPE.

## Introduction

Degeneration of rod and cone photoreceptors causes blindness in patients with retinitis pigmentosa and age-related macular degeneration. Proteins encoded by >100 distinct genes linked to these diseases have remarkably diverse biochemical activities. We hypothesize that defects in many of these genes create, either directly or indirectly, metabolic stress in photoreceptors. To address the validity of this idea, we are exploring photoreceptor energy metabolism and the metabolic interactions between photoreceptors and their environment.

Photoreceptors in the outer retina receive nutrients from blood that flows through the fenestrated choriocapillaris just inside the sclera of the eye. Between the photoreceptors and this blood supply there is a monolayer of cells called the retinal pigment epithelium (RPE). The basal and apical plasma membranes of the RPE cells are enriched with the glucose transporter GLUT1 (Takata et al., 1992). Glucose is delivered from circulation to the outer retina via these transporters (Swarup et al., 2019).

Isolated retinas break down glucose through aerobic glycolysis and primarily convert the carbon from glucose into lactate (Warburg, 1925). This process, as opposed to oxidation of glucose carbons to CO_2_ in mitochondria, classically occurs in tumor tissue and is referred to as the Warburg effect (Warburg et al., 1927). Of the cells in the retina, photoreceptors express the same isoforms of glycolytic enzymes that are in tumor cells (Lowry et al., 1961; Lindsay et al., 2014; Casson et al., 2016; Rajala et al., 2016; Rueda et al., 2016). Retinas without photoreceptors consume less glucose and produce less lactate than normal retinas (Graymore, 1960; Reading and Sorsby, 1962; Winkler et al., 2004). These data suggest that aerobic glycolysis is performed by photoreceptors.

In a previous report, we described evidence that RPE cells minimize their consumption of glucose so that more glucose can traverse the RPE to reach the retina (Kanow et al., 2017). We also proposed that lactate produced by photoreceptors can suppress consumption of glucose by RPE cells (Kanow et al., 2017). That proposal was based on the results of *ex vivo* analyses of metabolic flux in retinas, RPE-choroid and cultured RPE cells. Here we tested our proposal more rigorously by using multiple unique mouse models, and by infusing ^13^C-labeled fuels into mice and quantifying ^13^C-labeled metabolites in their retinas and RPE-choroid tissues. We show that the retina primarily consumes glucose, that photoreceptors are the main consumers of glucose in the retina, and that lactate, both from circulation and from photoreceptors influences energy metabolism in the RPE.

## Results

### Retina metabolism is fueled primarily by circulating glucose

Rapid consumption of glucose and production of lactate are key metabolic features of isolated retinas (Warburg, 1925; Winkler, 1981; Ames et al., 1992). An isolated mouse retina incubated in culture medium produces nearly two moles of lactate from each mole of glucose that it consumes (Wang et al., 1997; Hass et al., 2023a). Isolated retinas can oxidize fuels other than glucose (Keen and Chlouverakis, 1965; Winkler, 1981; Chertov et al., 2011; Adijanto et al., 2014; Joyal et al., 2016; Kanow et al., 2017), yet the degree to these fuels are normally used *in vivo* is unknown.

We designed an experiment to compare consumption of several types of fuels by retinas within the eyes of awake, freely moving mice (Hui et al., 2017, 2020). Mice were equipped with jugular vein and carotid artery catheters. We infused glucose fully labeled with ^13^C (“^13^C_6_-glucose” or “m+6 glucose”) through the jugular veins. For comparison, in separate experiments we infused ^13^C_3_-lactate, ^13^C_4_-succinate, ^13^C_5_-glutamine, or ^13^C_4_-malate into jugular vein catheters for four hours (glucose) or two hours (each other fuel). Metabolite concentrations and infusion rates are listed in **Table 1**. **Fig. 1A** shows the experiment scheme, and **Fig. 1B** shows how these fuels integrate into energy metabolism. During the infusion we collected plasma from a carotid artery catheter every 30 minutes. Two or four hours after initiating the infusion, we collected final plasma samples, euthanized each mouse, dissected tissue samples, and froze them in liquid N_2_. Isotopic labeling of metabolites in plasma and tissue samples was quantified either by gas chromatography/mass spectrometry (GC-MS) or liquid chromatography/mass spectrometry (LC-MS/MS).

**Figure 1.**
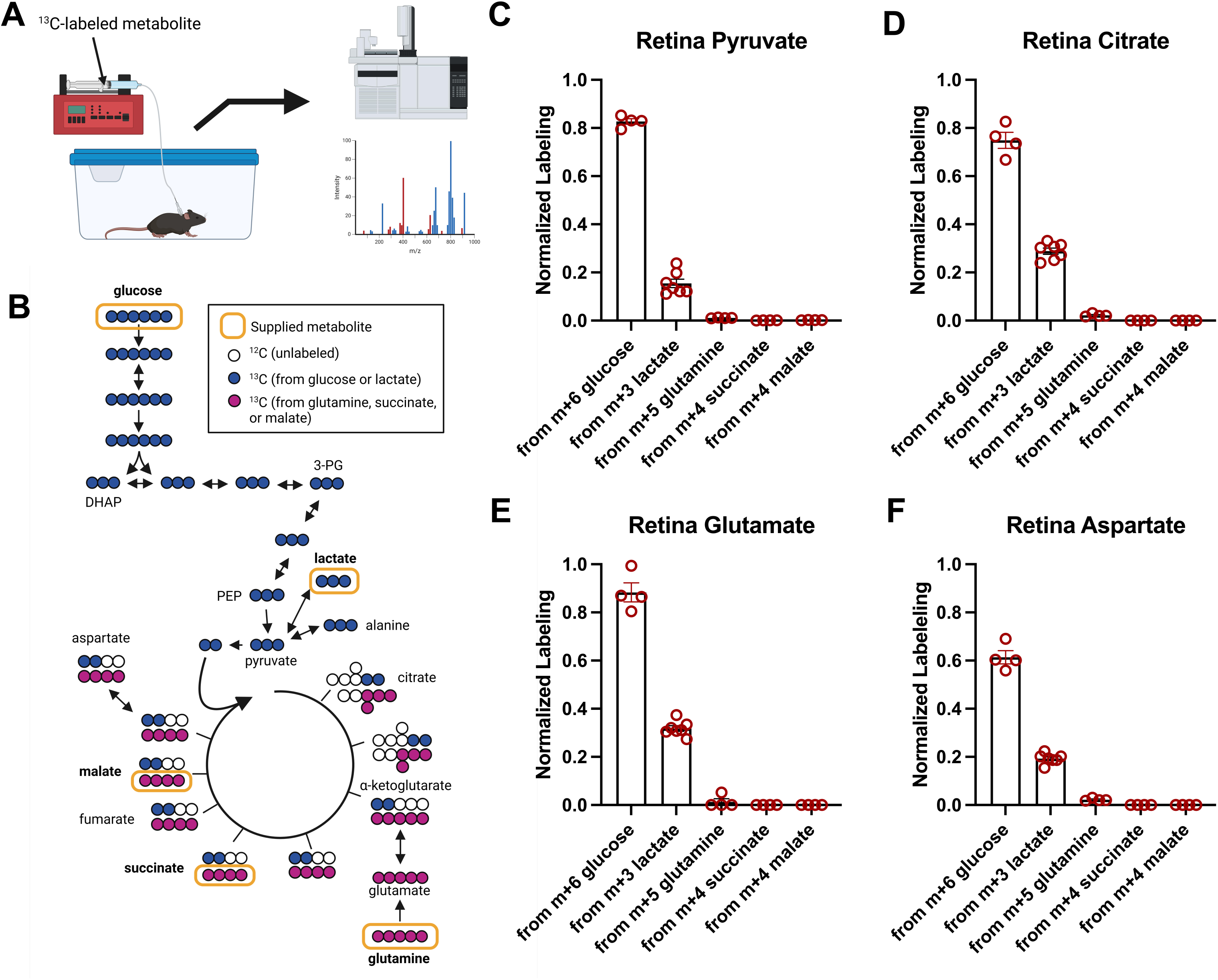
Glucose is the primary fuel for retina metabolism. Jugular vein and carotid artery catheters were installed in C57BL/6J mice. After mice had recovered from the surgery, they were infused with ^13^C_6_-glucose, ^13^C_3_-lactate, ^13^C_5_-glutamine, ^13^C_4_-succinate, and ^13^C_4_-malate. Infusion parameters are listed in **Table 1**. Shown in the figure are schematics of (**A**) the setup for infusion experiments, and (**B**) the pathways through which infused fuels (highlighted by yellow ovals) integrate into energy metabolism. Each circle represents a carbon atom on an intermediate in these pathways. Blue circles represent labeling patterns from glucose or lactate, while purple circles represent labeling patterns from glutamine, succinate, and malate. After mice are infused with each tracer, they are euthanized and their retinas dissected then. Tissue and plasma samples are analyzed by mass spectrometry. **C-F** show the labeling of (**C**) pyruvate, (**D**) citrate, (**E**) glutamate, and (**F**) aspartate from infused ^13^C_6_-glucose (n=4), ^13^C_3_- lactate (n=7), ^13^C_5_-glutamine (n=4), ^13^C_4_-succinate (n=4) or ^13^C_4_-malate (n=4). “Normalized labeling” indicates the labeling of the metabolic intermediate in the figure panel divided by the labeling of the infused intermediate in circulation. This normalization method accounts for differences in the proportion of circulating fuel in the bloodstream.

**Table 1.** Stock concentrations and infusion rates for ^13^C-labeled fuels.

| Metabolite | Concentration (mM) | Infusion rate (μL/g/min) | Infusion period (min) | Supplier | Catalog # |
| --- | --- | --- | --- | --- | --- |
| <sup>13</sup>C6-glucose | 200 | 0.1 | 240 | Cambridge Isotope Laboratories | CLM-504 |
| <sup>13</sup>C3-lactate | 400 | 0.1 | 120 |  | CLM-1579 |
| <sup>13</sup>C4-succinate | 10 | 0.05 | 120 |  | CLM-1571 |
| <sup>13</sup>C5-glutamine | 100 | 0.1 | 120 |  | CLM-1822-H |
| <sup>13</sup>C4-malate | 10 | 0.05 | 120 |  | CLM-8065-PK |

Plasma samples were used to confirm that each infused fuel reached isotopic steady state in circulation (**Supplementary Fig. 1A-E**). At isotopic steady state, we determined the extent to which each infused fuel is used to make m+3 pyruvate (**Fig. 1C**), m+2 citrate (**Fig. 1D**), m+2 glutamate (**Fig. 1E**), and m+2 aspartate (**Fig. 1F**) in the neural retina. The y-axis for these figure panels shows normalized labeling, which is calculated as the fractional enrichment of a metabolite (including all isotopologues) with ^13^C in the tissue relative to the fractional enrichment of the source of ^13^C in plasma (Hui et al., 2017, 2020). When the infused fuel is ^13^C_6_-glucose, a normalized labeling value for pyruvate of 1.0 would mean that 100% of pyruvate produced in the tissue was produced from glucose that had been taken up from circulating blood.

Metabolic intermediates in the retina are primarily derived from ^13^C_6_-glucose and ^13^C_3_-lactate, and poorly labeled from all other sources. m+6 glucose generates circulating m+3 lactate (**Supplementary Fig. 2A**), and carbons from circulating m+3 lactate can contribute to ^13^C- labeled pyruvate, citrate, glutamate, and malate in the retina (**Fig. 1C-F**). Therefore, a portion of glucose-derived labeling can occur via lactate (Hui et al., 2017). Similarly, ^13^C-lactate can undergo gluconeogenesis and generate circulating ^13^C-labeled glucose (**Supplementary Fig. 2B**) (TeSlaa et al., 2021). We calculated the direct contributions of glucose or lactate to the TCA cycle (approximated by averaging ^13^C labeling on citrate, α-ketoglutarate, glutamate, and aspartate) in the retina using a previously described formula (Described in Supplementary note 3 of (Hui et al., 2017)). This type of analysis produces an unrealistic result, that 114% of the retina’s TCA cycle is directly fueled by glucose, and 8.5% by lactate. However, this calculation is impacted by m+1 labeling on TCA cycle intermediates, which is abundant (pink bars in **Supplementary Fig. 2C**). M+1 labeling may be an artifact of ^13^CO_2_ that was generated by other tissues but recycled in the retina (Duan et al., 2022). When m+1 isotopologues are not included, the revised calculation indicates that glucose and lactate fuel 73.9% and 3.3% of the retina’s TCA cycle (**Supplementary Fig. 2D**). The primary message of these calculations is that glucose is the major TCA cycle fuel in the retina, and that the direct contribution of circulating lactate to retina metabolism is small.

### Photoreceptors are the main consumers of glucose in the retina

Previous studies showed that glucose consumption increases while rod photoreceptors are developing within the retina, and decreases when photoreceptors degenerate (Cohen and Noell, 1960; Reading and Sorsby, 1962). We compared rates of glucose consumption by normal retinas (C57BL/6J) vs. by retinas without photoreceptors (*Aipl1*^-/-^ in the C57BL/6J background). *Aipl1*^-/-^ mice are models of Leber congenital amaurosis. In these mice, rod and cone photoreceptors degenerate fully by one month of age (Ramamurthy et al., 2004). We used this model to confirm the contribution of photoreceptors to retina glucose consumption.

We used an enzymatic assay to quantify depletion of glucose from the medium bathing freshly isolated retinas. *Aipl1*^-/-^ retinas deplete glucose from the medium at only 15% of the rate of C57BL/6J controls and they release lactate at 22% of the control rate (**Fig. 2A-B**). We also measured accumulation of ^13^C_3_ lactate within the isolated retinas when they were incubated in ^13^C_6_-glucose. The initial rate of ^13^C lactate accumulation within *Aipl1*^-/-^ retinas is only 11% of the rate of C57BL/6J controls (**Fig. 2C**).

**Figure 2.**
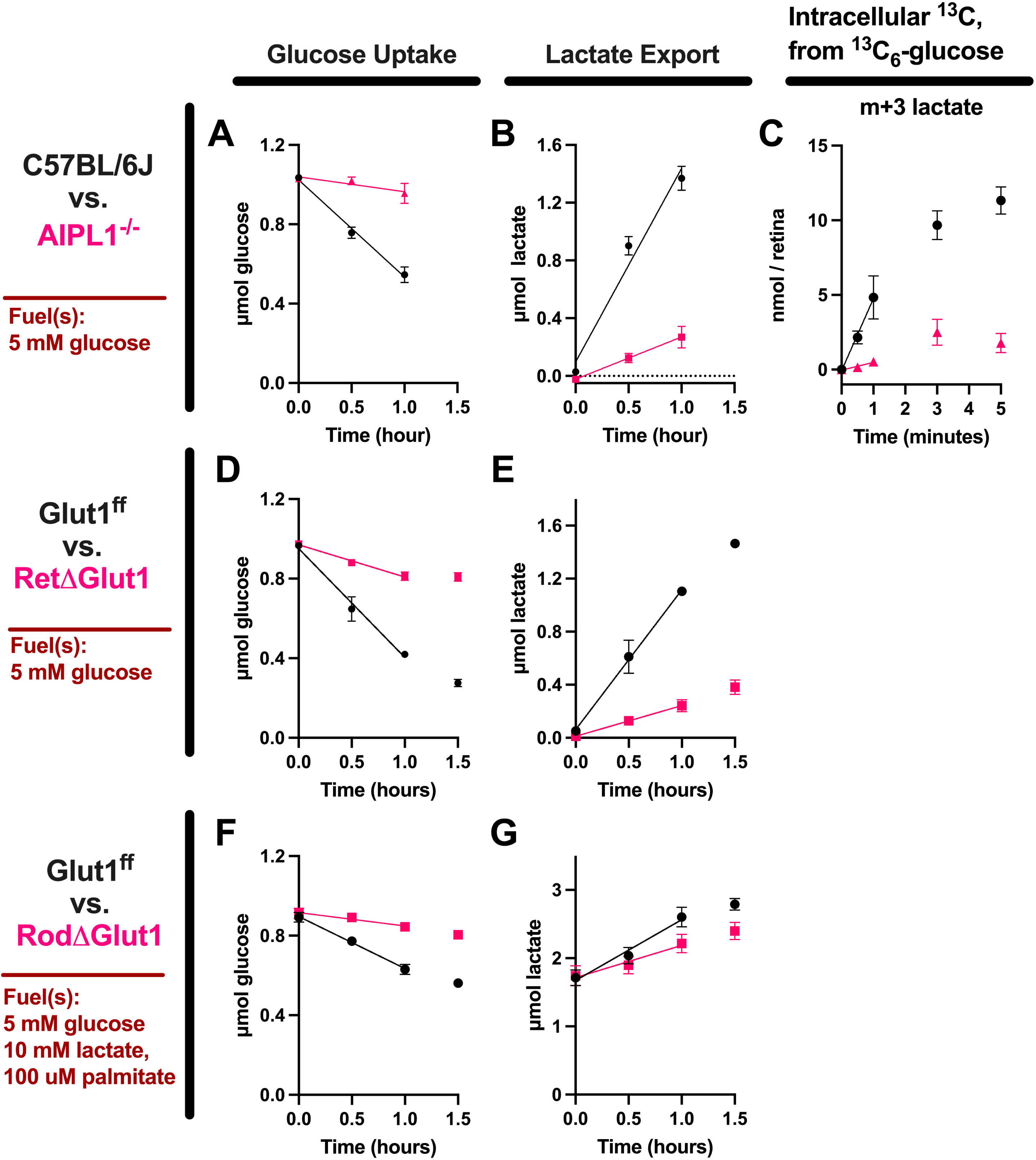
Photoreceptors are the main consumers of glucose in the retina. Retinas were dissected from eyes then incubated in Krebs-Ringer Bicarbonate buffer supplemented with 5 mM glucose, at 37°C and 5% CO_2_. Glucose consumption from medium (**A**, **D**, **F**) and lactate production (**B**, **E**, **G**) were monitored over 1-1.5 hours. These assays were used to determine how glycolytic metabolism is affected by (**A**, **B**) photoreceptor degeneration due to loss of AIPL1 (n=3-4), (**D**, **E**) loss of Glut1 from the retina (n=4-5), or (**F**, **G**) loss of Glut1 from rod photoreceptors (n=5). Loss of photoreceptors, loss of retinal Glut1, or loss of rod Glut1 slowed retina glucose utilization. (**C**) C57BL/6J and AIPL1^-/-^ retinas were also incubated in KRB supplemented with 5 mM ^13^C_6_-glucose. Retina tissue was flash-frozen 0, 0.5, 1, 3, or 5 minutes after the start of the incubation. Labeling of metabolic intermediates in the retina was assessed by mass spectrometry. Labeling of lactate with ^13^C from glucose is slowed by the loss of photoreceptors (n=3-4 retinas / time point).

GLUT1 is the only glucose transporter that normally is expressed in detectable amounts in rods (Daniele et al., 2022). If photoreceptors consume most of the glucose provided to retinas, then deletion of Glut1 from photoreceptors should diminish glucose consumption substantially. We determined initial rates of glucose depletion and lactate accumulation in culture medium by retinas from mice in which *Glut1* was conditionally deleted either from the whole retina (RetΔ*Glut1*) or from rods only (RodΔ*Glut1*). Loss of *Glut1* from the entire retina slows the rate of glucose depletion to 30% of the rate of Glut1^fl/fl^ control retinas (**Fig. 2D**) and lactate export to 22% (**Fig. 2E**). In the experiment comparing RodΔ*Glut1* mice to controls, glucose consumption and lactate production were assessed in the presence of 10 mM lactate and 100 µM palmitate-BSA in the KRB buffer bathing retinas. Loss of *Glut1* from rod photoreceptors slows glucose depletion to 26% of *Glut1^fl^*^/fl^ controls **Fig. 2D**) and lactate export to 53% (**Fig. 2E**). These data confirm that photoreceptors are the primary cell type that takes up glucose and exports lactate, and that they import glucose mostly through GLUT1.

### Photoreceptors can use lactate for fuel when glucose is unavailable

Remarkably, despite being the main consumers of glucose in the retina, rod photoreceptors do not die immediately upon loss of components essential for glycolysis or for uptake of glucose (Chinchore et al., 2017; Daniele et al., 2022; Weh et al., 2023). That suggests that rods can use other fuels when glucose is not available. We hypothesized that under normal conditions oxidation of glucose by a rod consumes NAD^+^, so its capacity to import and oxidize lactate is limited. But when glucose is not available for oxidation the cytosolic NAD^+^/NADH ratio in rod should increase. A greater abundance of cytosolic NAD^+^ could promote oxidation of cytosolic lactate to pyruvate. This would deplete intracellular lactate and generate a gradient of lactate that favors import of extracellular lactate (**Fig 3A**).

**Figure 3.**
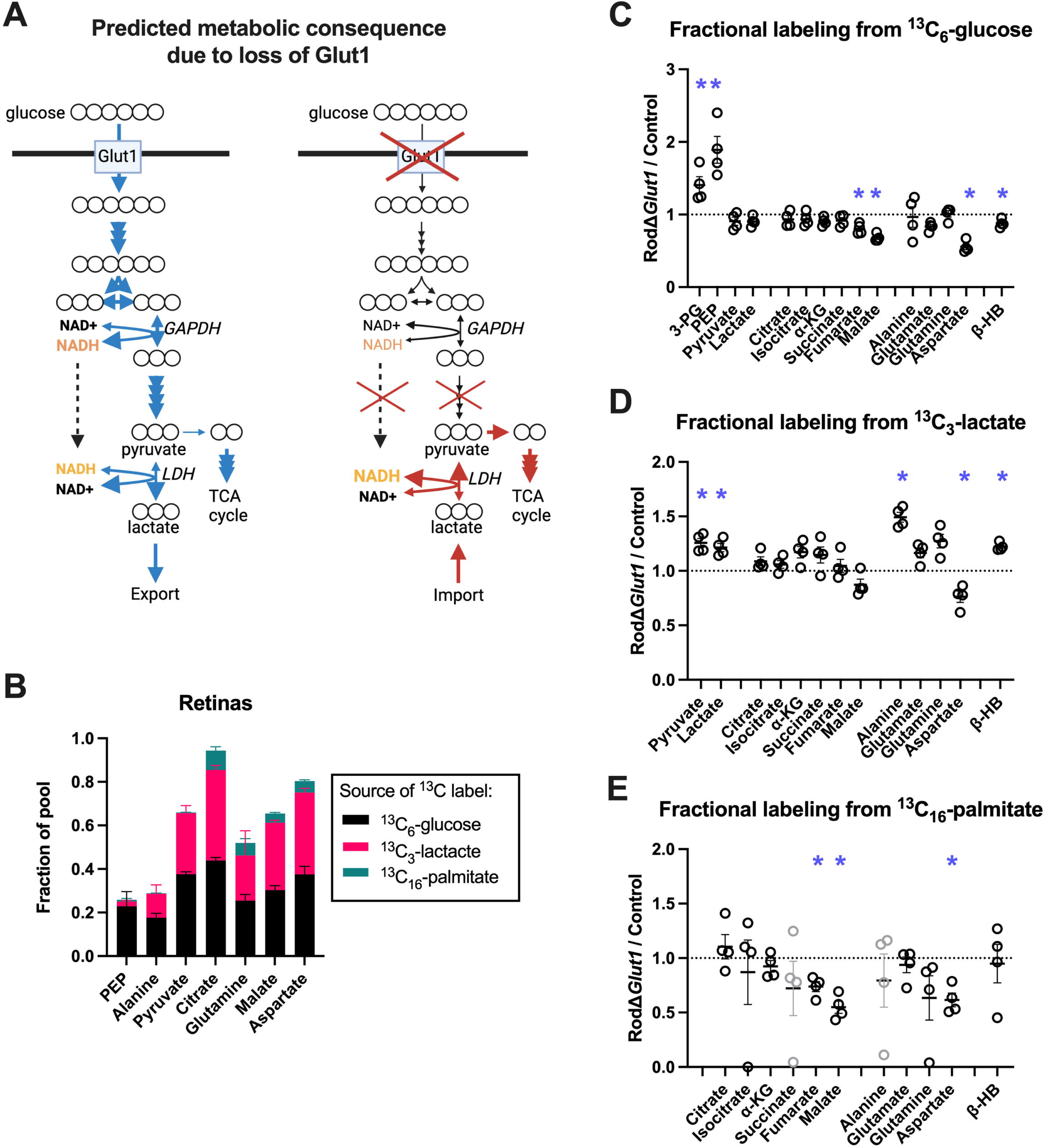
When retinas cannot consume glucose, they consume more lactate. Mice lacking Glut1 in rod photoreceptors (RodΔGlut1) and identically treated littermate control mice were euthanized, and retinas dissected. Retinas were incubated for 2 hours in KRB supplemented with 5 mM glucose, 10 mM lactate, or 100 µM palmitate-BSA. In this experiment, source of ^13^C was ^13^C_6_-glucose for one cohort of retinas, ^13^C_3_-lactate for the second cohort, or ^13^C_16_-palmitate in a third cohort. Following the incubation, retinas were flash-frozen in liquid N_2_ and later analyzed by mass spectrometry. Our hypothesis is that loss of glucose metabolism in rods would increase metabolism from the other fuel sources by altering NAD^+^/NADH. This hypothesis is illustrated in (**A**). (**B**) in control retinas, the level of labeling on PEP, alanine, pyruvate, citrate, glutamine, malate, or aspartate from ^13^C_6_-glucose, ^13^C_3_-lactate, or ^13^C_16_-palmitate. Glucose and lactate are the most prevalent sources of ^13^C. Fractional labeling of glycolytic, TCA cycle intermediates, and amino acids in control and RodΔGlut1 retinas from (**C**) ^13^C_6_-glucose (n=3-4), (**D**) ^13^C_3_-lactate (n=3-4), or (**E**) ^13^C_16_-palmitate (n=4). Data are displayed as a fold-change from labeling in control retinas.

We compared fuel preferences of isolated mouse retinas in KRB supplemented with 5 mM glucose, 10 mM lactate, and 100 µM palmitate-BSA. We included palmitate because previous reports had shown that it can be a respiratory fuel for retinas (Joyal et al., 2016). We used three treatment conditions. The overall concentration of each metabolic fuel remained the same across treatment conditions, but in each treatment condition only one of the fuels was labeled with ^13^C. We isolated retinas from control and RodΔGlut1 mice and then incubated them for 2 hours in these media. We harvested the retinas, extracted metabolites from them, and used GC-MS to quantify how the carbons from each type of fuel had integrated into metabolic intermediates. ^13^C from each fuel can integrate its carbons into TCA cycle intermediates, but glucose and lactate substantially outperformed palmitate under these conditions (**Fig. 3B**).

We compared consumption of fuels by control and RodΔGlut1 retinas. We tested the idea that rods oxidize other fuels when access to glucose is decreased. The absence of GLUT1 in rods substantially influences the abundance of metabolic intermediates in the retina (**Supplementary Fig. 3**), and how well these intermediates were ^13^C labeled from each fuel (**Fig. 3C-E**).

Fractional labeling from ^13^C glucose decreased for most metabolic intermediates, suggesting a diminished ability of glucose to fuel a portion of retina metabolism (**Fig. 3C**). However, fractional labeling of the glycolytic intermediates 3-PG and PEP increased substantially in the absence of GLUT1 in rods. Notably, it has been reported that Müller cells express low levels of pyruvate kinase (PK) (Lindsay et al., 2014). Abnormally high uptake of ^13^C glucose into Müller cells could cause a buildup of ^13^C labeled 3PG and PEP upstream of the PK reaction. A similar finding for *Glut1*-deficient macrophages has been reported (Freemerman et al., 2019).

The absence of GLUT1 in rods limits incorporation of ^13^C from glucose, but incorporation of ^13^C from lactate increases (**Fig. 3D**). That finding suggests that lactate consumption increases to compensate for the diminished ability to import glucose into rods. Pyruvate and several other intermediates were labeled more in RodΔ*Glut1* retinas by ^13^C_3_-lactate. Incorporation of carbons from ^13^C_16_-palmitate decreased in RodΔ*Glut1* retinas, suggesting that glucose and palmitate metabolism in retinas are synergistic (**Fig. 3E**). Our findings suggest that when the supply of glucose to photoreceptors is cut off, photoreceptors instead catabolize lactate.

### *In vivo* evidence that RPE imports lactate that has been exported from photoreceptors

Normally, photoreceptors are the primary exporters of lactate in mouse retinas. **Figs. 2B** and **2C** show that when fueled with glucose, *Aipl1*^-/-^ retinas export much less lactate than normal retinas. Several groups have demonstrated that RPE can transport lactate, and that lactate RPE transport is essential for retinal health (la Cour et al., 1994; Daniele et al., 2008). Kanow et al. proposed that lactate that has been exported from the retina is metabolized within the RPE (Kanow et al., 2017). That hypothesis was based on findings from *ex vivo* analyses of tissues and cultured RPE cells. We designed an *in vivo* experiment that uses retinas without photoreceptors to test this hypothesis in the eyes of living mice. We predicted that without photoreceptors to export lactate, less lactate will be imported into the RPE-choroid.

We dissected RPE-choroid tissue from control mice and from two different lines of mice in which photoreceptors had degenerated, *Aipl1*^-/-^ and *Pde6b^H620Q^*^/H620Q^. *Pde6b^H620Q^*^/H620Q^ mice are models of retinitis pigmentosa, with a slower form of retinal degeneration than *Aipl1*^-/-^ mice (Davis et al., 2008a). We used GC-MS to quantify the abundance of metabolites in RPE-choroid tissue of these models (**Fig. 4A-F**). Lactate is the only glycolytic metabolite in the RPE-choroid that consistently was less abundant when photoreceptors in the retina had degenerated (**Fig. 4A,D**). This finding further supports the idea that normally the RPE imports lactate made by and released from photoreceptors (Daniele et al., 2008).

**Figure 4.**
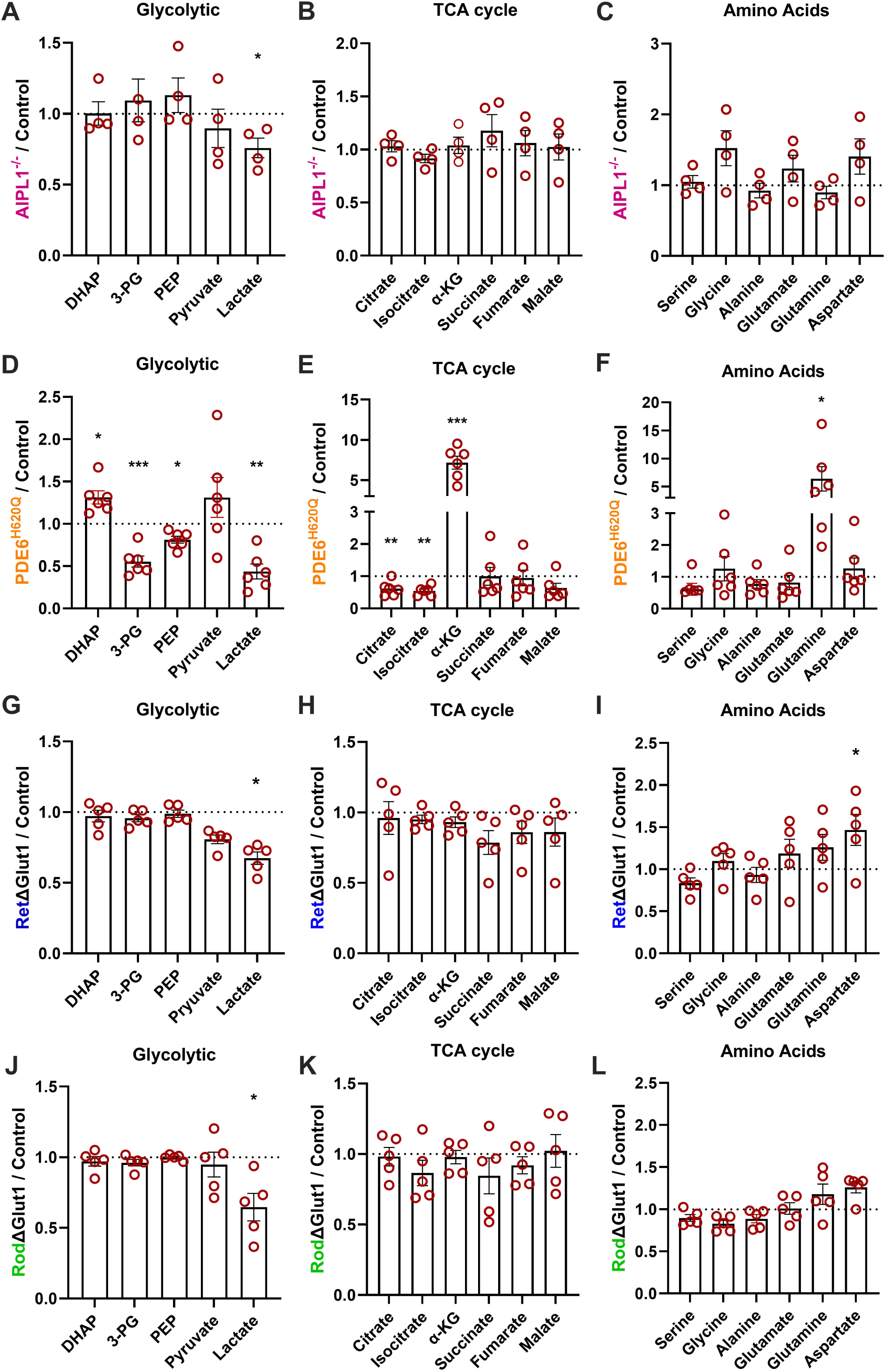
When photoreceptors do not consume glucose, there is a decrease in RPE- choroid lactate. *Aipl1*^-/-^, *Pde6b^H620Q^*, RetΔGlut1, RodΔGlut1, and control mice were euthanized and RPE-choroid tissue was immediately dissected from the eye and flash-frozen. Tissue metabolite levels were analyzed using mass spectrometry, and divided into three categories; glycolytic intermediates (**A, D, G, J**), TCA cycle intermediates (**B, E, H, K**), and amino acids (**C, F, I, L**). Metabolite abundances in *Aipl1*^-/-^ (**A-C**) and *Pde6b^H620Q^*RPE-choroid tissue (**D-F**) was compared to metabolite abundances in RPE-choroid tissue from C57BL/6J controls of a similar age. Metabolite abundances in RetΔGlut1 (**G-I**) and RodΔGlut1 RPE- choroid tissue (**J-L**) was compared to metabolite abundances in RPE-choroid tissue from uninjected or tamoxifen injected Glut1^fl/fl^ controls of a similar age. Lactate levels were consistently lower in the experimental groups of all tissues. n=5-6 per group.

We also tested this idea by quantifying metabolites in RPE-choroid from mice in which GLUT1 either is absent from the entire retina (<u>Ret</u>Δ*Glut1*), or absent only from rods (<u>Rod</u>Δ*Glut1*). As with the models of photoreceptor degeneration, there is less lactate in the RPE-choroid from eyes of these mice (**Fig. 4G-L**). Lactate was not the only metabolite with altered levels in these models, but it was the only metabolite that was altered consistently in the RPE-choroid of all four model systems. In three of four models, there also was an increase in aspartate, suggesting either an increase in retina-derived aspartate, an increase in local aspartate production, or a decrease in local aspartate consumption. We also noted a substantial increase in α-ketoglutarate and glutamine uniquely in the PDE6-deficient RPE-choroid tissue.

### *In vivo* flux measurements confirm that lactate produced by photoreceptors serves as an energetic fuel for RPE-choroid

The next set of experiments show that AIPL1-deficiency directly affects metabolic flux from glucose only in retina tissue and not in RPE tissue.

*Ex vivo analysis*: *Aipl1* normally is expressed in photoreceptors and not in RPE cells (van der Spuy et al., 2002). There are no photoreceptors in adult *Aipl1*^-/-^ mouse retinas because they all have degenerated. We isolated retina and RPE-choroid tissue from C57BL/6J and *Aipl1*^-/-^ mice, incubated them separately in 5 mM ^13^C_6_-glucose and harvested tissues between 0 and 5 minutes after beginning the incubation. **Fig. 5A** shows that AIPL1^-/-^ retinas produce much less lactate than control C57BL/6J retinas. **Fig. 5B** shows that lactate production is not altered in RPE-choroid tissue isolated from *Aipl1* ^-/-^ mice. Normally, retinal tissue produces much more lactate than RPE-choroid tissue. The slower production of lactate that does occur in the RPE-choroid is not affected by loss of *Aipl1*.

**Figure 5.**
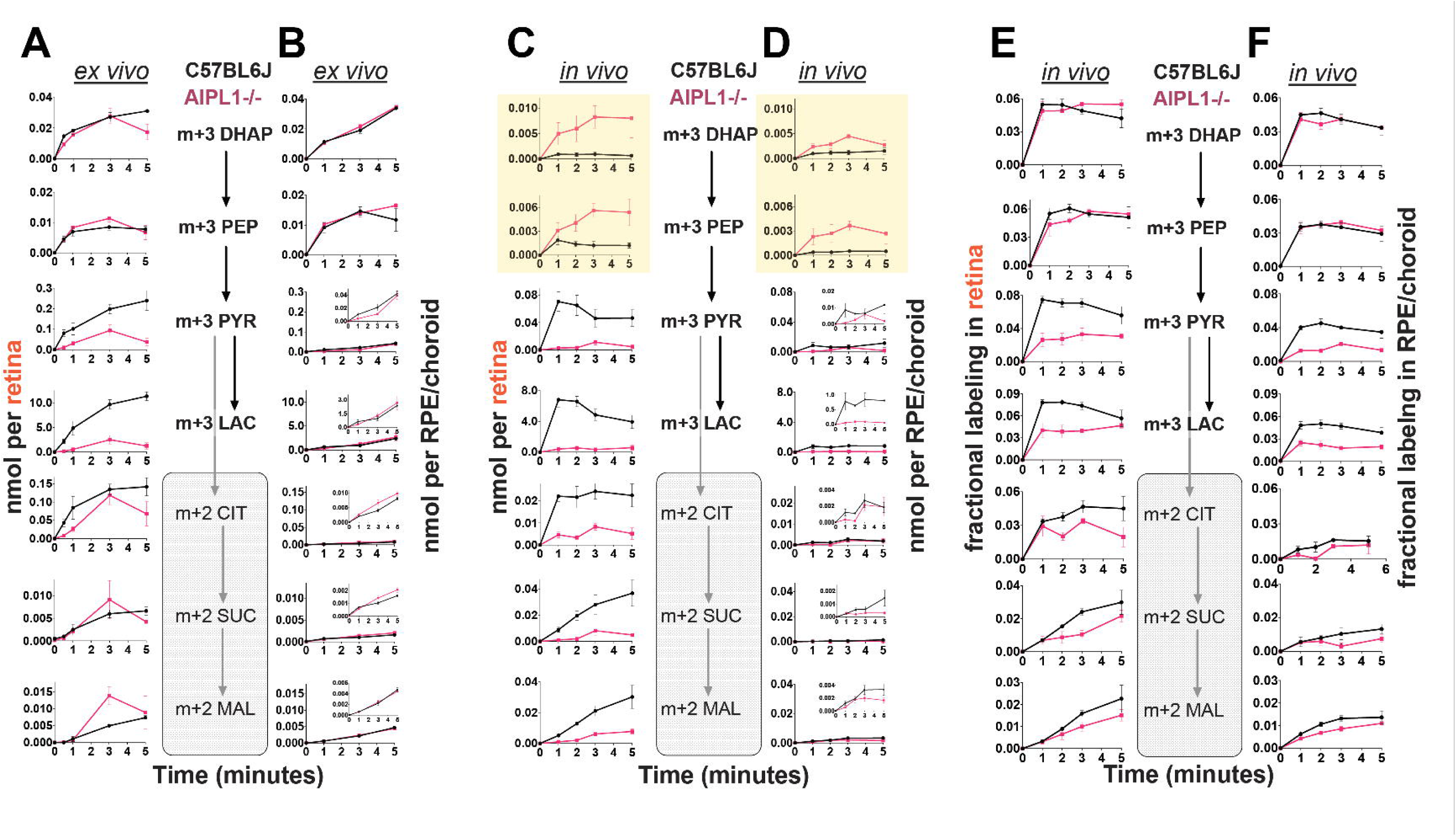
Mouse RPE-choroid uses photoreceptor-derived lactate *in vivo.* (A, **B)** C57BL/6J and *Aipl1* ^-/-^ mice were euthanized and their (**A**) retina or (**B**) RPE-choroid tissue were incubated *ex vivo* in ^13^C_6_-glucose. The amount of ^13^C labeled DHAP, PEP, pyruvate, lactate, citrate, succinate, and malate were quantified by mass spectrometry. *Aipl1*^-/-^ decreases utilization of glucose by the retina, but not the RPE-choroid. (**C, D**) Jugular vein catheters were installed in C57BL/6J and *Aipl1*^-/-^ mice. A bolus dose of 100 mg/kg ^13^C_6_-glucose was infused through jugular vein catheters, then mice were euthanized 1, 2, 3, or 5 minutes later. Tissue levels and ^13^C labeling of DHAP, PEP, pyruvate, lactate, citrate, succinate, and malate were determined using mass spectrometry. ^13^C labeling of labeling of metabolic intermediates was drastically altered in both the (**C**) retina and (**D**) RPE-choroid of mice where the ^13^C_6_-glucose had been infused. (**E, F**) contain data from the same mice as **C** and **D**, but show the proportion of each metabolite that is ^13^C labeled. In all panels, “0 minute” samples were from retina or RPE-choroid tissue that was not exposed to a ^13^C fuel.

Those results confirm that photoreceptors must be present for glycolysis to be the dominant metabolic pathway in the retina. AIPL1^-/-^ retinas do not make enough lactate to export it in abundance to the RPE.

*In vivo analysis:* We hypothesized that lactate imported from photoreceptors normally is a significant fraction of the total lactate in RPE-choroid tissue. Disrupted delivery of glucose-derived lactate from the retina to the RPE would limit the amount of lactate in RPE-choroid. We designed an experiment to test this.

We infused a bolus dose of 100 mg/kg ^13^C_6_-glucose into jugular vein catheters in C57BL/6J and *Aipl1* ^-/-^ mice. We euthanized mice 1, 2, 3, or 5 minutes following the infusion, and dissected the retina and RPE-choroid. We extracted metabolites from each tissue and used mass spectrometry to quantify the extent to which carbons from ^13^C_6_-glucose were used to make glycolytic and mitochondrial intermediates (**Fig. 5C and D**). The result confirms our prediction that much less m+3 lactate reaches the RPE-choroid **(Fig. 5D)** in AIPL1^-/-^ mice than in controls.

Nevertheless, other findings from that *in vivo* experiment were unexpected and puzzling. We found that ^13^C labeled glycolytic intermediates (DHAP and PEP) upstream of the reaction catalyzed by PK seemed to accumulate both in retinas and in RPE-choroid from AIPL1^-/-^ eyes. That result appeared inconsistent with our *ex vivo* analyses (**Fig. 5A,B**) and our predictions.

Further analyses revealed the explanation for this unexpected observation. There is an unavoidable delay of ∼2 minutes between euthanasia and the time when retina and RPE-choroid tissues are fully dissected and frozen. During that interval there is no fresh blood available to the tissues to maintain a supply of glucose. We hypothesized that due to the rapid consumption of glucose by photoreceptors in normal retinas, postmortem metabolism could deplete glycolytic intermediates rapidly in the tissue. Metabolites would be depleted faster in wild-type retinas that have photoreceptors than in AIPL1^-/-^ retinas during the time between euthanasia and tissue harvest.

We performed an *ex vivo* experiment to test this hypothesis. We incubated isolated retinas from C57BL/6J and AIPL1^-/-^ mice for 10 minutes in KRB medium supplemented with 5 mM ^13^C_6_- glucose. Then we moved the retinas to KRB medium without glucose. We collected retinas at 0, 1, 3, and 5 minutes after moving the tissue, and then quantified glycolytic intermediates.

Remarkably, the absolute quantities of ^13^C-labeled DHAP, 3PG and PEP (black lines in **Supplementary Fig. 4A**) in normal retinas decreased rapidly, 50-70% within the first minute. In contrast, these intermediates were stable in AIPL1^-/-^ retinas (red lines in **Supplementary Fig. 4A)**. We also monitored the fractional labeling of the metabolites (**Supplementary Fig. 4B**). The fractional labeling of early glycolytic intermediates also decreased within the first minute.

However, the fractional labeling of later intermediates, pyruvate and lactate remained closer to their initial values at the time that glucose was withdrawn.

This result of this *ex vivo* experiment is consistent with our hypothesis. What had appeared to be accumulation of DHAP and PEP in AIPL1^-/-^ tissues relative to control tissues (**Fig. 5C)** is caused by an artifact. The artifact occurs during the unavoidable delay between euthanasia and harvesting of the tissue. It is caused by very rapid consumption of glucose by photoreceptors in control retinas. That rapid consumption does not occur in AIPL1^-/-^ retinas. The same effect occurs in RPE-choroid tissue after blood flow to the eye has stopped. During that time the wild-type retina, which is in close contact with the RPE, can consume glucose rapidly from the RPE until we separate the tissues in the final step of the dissection.

The *ex vivo* simulation shows that fractional labeling of DHAP and PEP at the time of euthanasia are not well preserved during 1-2 minutes of ischemia (**Supplementary Fig. 4B**). However, fractional labeling of pyruvate and lactate are better preserved. **Fig. 5E and F** show the results of the same bolus infusion experiment as in **Fig. 5C and D** but instead of the data being expressed as absolute amounts they are expressed as fractional labeling. It shows that pyruvate and lactate are substantially less labeled both in the retina and in the RPE of mice that lack photoreceptors in their retinas. Even though glycolytic intermediates are depleted faster the RPE-choroid from control mice still have as much as 9 times more lactate **(Fig. 5D)** than the RPE-choroid from eyes that do not have photoreceptors.

In summary, the *in vivo* bolus infusion confirms that less lactate reaches the RPE-choroid when there are no photoreceptors in the retina.

## Discussion

Normal vision is dependent on the circulation of carbon between the retina and RPE (Daniele et al., 2008, 2022). The retina and the RPE provide metabolic support for each other (Lakkaraju et al., 2020). Energy from light isomerizes 11-cis retinal to all-trans in the retina (Wald, 1968). The visual cycle in the RPE then uses metabolic energy to regenerate 11-cis retinal for the retina (Saari, 2016). Photoreceptors also use substantial carbon to generate outer segments disks, which constantly turn over and which require glucose transport (Young, 1967; Swarup et al., 2019; Daniele et al., 2022). The RPE also recycles material from the tips of rod outer segments, and it transports ions, nutrients and water to and from the retina (Young and Bok, 1969; Quinn and Miller, 1992; la Cour et al., 1994). In this report we focused on energy metabolism.

In a previous report, we described distinct metabolic features of the retina and RPE and proposed that they are complementary and optimized to ensure that sufficient glucose can traverse the RPE to fuel the retina (Kanow et al., 2017). That hypothesis was based primarily on biochemical measurements from isolated mouse retina tissue, mouse RPE-choroid tissue and cultured human RPE cells. After proposing the hypothesis, which we referred to as a “metabolic ecosystem”, we felt that a more direct demonstration of metabolic relationships in the eye of a living animal would better support this hypothesis. This report describes those tests and several new observations.

### We extended our previous findings to show that

- Retina metabolism in the eye of a living mouse is fueled primarily by circulating glucose and lactate
- Photoreceptors are the main consumers of glucose in the retina
- Photoreceptors can use lactate for fuel when glucose is unavailable
- RPE tissue in the eye of a living mouse imports lactate that has been exported from photoreceptors
- Lactate produced by photoreceptors in the eye of a living mouse can be oxidized to TCA cycle intermediates by RPE-choroid tissue

### Why do photoreceptors consume so much glucose?

Photoreceptors hydrolyze ATP to maintain an essential ion gradient across their plasma membrane, and they use the energy extracted from glycolysis to synthesize new ATP (Ames et al., 1992; Okawa et al., 2008; Ingram et al., 2020). We show that most of the glycolytic activity in the retina within the eye of a living mouse happens in photoreceptors. Because photoreceptor and RPE mitochondria consume so much O_2_, the steady state concentration of O_2_ in the outer retina is very low (Linsenmeier, 1986; Yu and Cringle, 2006). In any biological system, mitochondrial activity is limited by O_2_ availability, and glycolysis necessarily becomes a predominant metabolic pathway from which to extract energy from fuels (Gstraunthaler et al., 1999; Gilglioni et al., 2018; Hass et al., 2023b; Tan et al., 2024).

It is possible that retinas benefit from limited availability of O_2_. Along with light, O_2_ is a major source of retina lipid peroxidation (Wiegand et al., 1983). Maintaining low O_2_ levels in the retina could slow or prevent lipid peroxidation. A highly active mitochondrial metabolism in photoreceptors and RPE could therefore help to shield retina lipids from peroxidation.

Glucose also is used in anabolic pathways, and it is necessary for outer segment synthesis (Daniele et al., 2022). It is likely that a far greater proportion of glucose turnover feeds into catabolic pathways than into anabolic pathways. Yet the absence of glucose catabolism does not result in immediate photoreceptor death (Chinchore et al., 2017; Daniele et al., 2022; Weh et al., 2023), and to an extent, lactate can substitute for glucose as an energetic fuel. This may imply that if there is a defect that impacts glycolysis, increasing lactate supply or catabolism could be a therapeutic mechanism to sustain photoreceptors.

### What sustains the RPE?

We showed in this report that in the eye of a living animal lactate exported from the retina can be imported into RPE and oxidized in mitochondria. Lactate slows the consumption of glucose in RPE cells and mouse RPE-choroid (Kanow et al., 2017). Lactate may therefore protect glucose from being consumed by the RPE, and allow for more glucose to fuel photoreceptor metabolic activity (Kanow et al., 2017). However, lactate is not the only fuel that RPE cells consume. They metabolize proline (Zhu et al., 2023), succinate (Bisbach et al., 2020; Hass et al., 2022), glutamate (Xu et al., 2020), and fatty acids (Adijanto et al., 2014; Reyes-Reveles et al., 2017). RPE cells also have the ability to store and mobilize glycogen (Senanayake et al., 2006). It will be important to establish the relative contributions of each of these fuels to RPE metabolism in the eye of a living animal and to determine how they are influenced by illumination, circadian factors, aging and disease.

## Acknowledgements

We thank Craig Beight for maintaining the colony of Rodι1Glut1 mice and for injecting tamoxifen.

## Funding Support

DTH is supported by a Brightfocus Foundation Postdoctoral Fellowship (M2022003F) and NEI K99EY034881. JYSH is supported by a Brightfocus Foundation Postdoctoral Fellowship (M2024001F). CMB was supported by F31EY031165. NDN is supported by F31EY033660.

SHT is supported by U01EY030580, U01EY034590 R24EY028758, R24EY027285, 5P30EY019007, R01EY033770, R01EY018213, R01EY024698, Piyada Phanaphat fund, Lynette & Richard Jaffe, NYEE Foundation, Foundation Fighting Blindness TA-GT-0321-0802- COLU-TRAP and RPB unrestricted grant. SHT is on the scientific and clinical advisory board for Emendo, Medical Excellence Capital, and Nanoscope Therapeutics. NSP is supported by P30EY015585 and IK6BX005233. NJP is supported by R01EY026525. JBH is supported by NEI R01EY06641, R01EY017863, R21032597 and Foundation Fighting Blindness TA-NMT- 0522-0826-UWA-TRAP. This work was supported in part by NIH NIDDK grant P30 DK017047 to the University of Washington Diabetes Research Center and used the Diabetes Research Center - Metabolic and Cellular Phenotyping Core.

## Materials and Methods

### Ethical approval

This study was carried out in accordance with the National Research Council’s Guide for the Care and Use of Laboratory Animals (*8th ed*). For experiments using C57BL6J and AIPL1^-/-^ mice, all protocols were approved by the Institutional Animal Care and Use Committees (IACUC) at the University of Washington and the ACORP at the Veterans Affairs – Puget Sound Health Care System. For experiments using tamoxifen-treated RodΔGlut1 mice and *Glut1^LoxP^*^/LoxP^ controls, protocols were approved by the IACUC at the Cole Eye Institute of the Cleveland Clinic. For experiments using RetΔGlut1 mice and Glut1^LoxP/LoxP^ controls, protocols were approved by the IACUC at the Thomas Jefferson University. For experiments using Pde6b^H620Q/H620Q^ mice, protocols were approved by the IACUC at Columbia University.

### Animals

All experiments adhered to our approved IACUC protocol and the ARVO statement for the use of animals in ophthalmology research. Mice were group housed at an ambient temperature of 25°C, with a 12-hour light cycle, and *ad libitum* access to water and normal rodent chow. We used male and female wild-type C57BL/6J (RRID: IMSR_JAX:000664) mice, and AIPL1^-/-^ mice bred to a C57BL/6J background over a series of at least six back-crosses. Mouse ages ranged from 2-6 months old except for RetΔGlut1, which were used at 1 month of age to avoid significant retinal degeneration. RodΔGlut1 mice were used at 2.5 months of age. We determined *ex vivo* glucose uptake, *ex vivo* lactate release, and metabolite content in tissue from Glut1^LoxP/LoxP^; crx-cre (RetΔGlut1) and *Glut1*^LoxP/LoxP^;*Pde6g-^creERT2^*(RodΔGlut1) mice. The generation of these transgenic mouse models is described in previous publications (Koch et al., 2015; Daniele et al., 2022). To induce cre recombinase activity in RodΔGlut1 mice, animals were given daily intraperitoneal injections of 100 μg tamoxifen / g body weight over 3 days.

Injection formulations were made by dissolving tamoxifen at 100 mg/ml in ethanol and diluting this solution 1:10 in corn oil. Injections were given beginning at 5 weeks of age and were also administered to littermate controls (Glut1^LoxP/LoxP^). *Pde6b^H620Q/H620Q^* mice were generated as previously described (Davis et al., 2008b) and kept as homozygous breeders.

### Euthanasia and tissue dissection

Mice were euthanized by awake cervical dislocation. The time of day for euthanasia was after 11 am, to avoid variability due to peak outer segment phagocytosis. We dissected eyes from the mouse into room temperature Hank’s buffered salt solution (Gibco, Cat#: 14170120). Eyes were cleared of extraocular muscle and collagenous tissue and the cornea was separated from the sclera by cutting along the ora serrata. We removed the cornea, iris, and lens, then teased apart the remaining retina and RPE-choroid tissue. This process typically takes ∼2-4 minutes for both eyes from one mouse. Retina and RPE-choroid tissue were immediately flash frozen in liquid N_2_ (*in vivo* experiments) or used in downstream experiments then flash frozen in liquid N_2_ (*ex vivo* metabolic flux).

### Installation of jugular vein and carotid artery catheters

The procedure for chronic jugular vein and carotid artery catheterization was performed as previously described (Ayala et al., 2011) by the Metabolic and Cellular Phenotyping Core of the University of Washington’s Diabetes Research Center. Briefly, following induction and maintenance under systemic inhaled anesthesia (isoflurane 1.5-2% in 1 mL/min), mice were administered with 4 mg/kg ketoprofen to reduce post-surgical swelling and provide analgesia. For intravenous infusions, jugular veins were isolated through a lateral incision to the trachea, and a silastic catheter was introduced into the vein, anchored to surrounding tissue, tunneled subcutaneously to the nape of the neck and connected to a vascular access port. A subset of mice received a contralateral carotid artery cannulation for blood sampling. The catheters were filled with heparinized saline, the skin incisions were sutured, and the mice recovered for 1 week before conscious infusion studies.

### Infusions

We infused labeled fuels one of two ways. In the first way (**Fig. 1**, **Supplemental Fig. 1**) jugular catheters were infused gradually through syringes connected to syringe pumps. Metabolites were dissolved in sterile saline and infused at the concentrations and rates listed in table 1.

In addition to long term infusions we administered 100 mg/kg ^13^C_6_-glucose as a bolus through jugular catheters, over a period of approximately 20 seconds. One, two, three, and five minutes after the start of the infusion, mice were euthanized. Retina and RPE-choroid tissue were dissected and snap frozen in liquid N_2_.

### Ex vivo metabolic flux

Unless otherwise specified, mouse retina and RPE-choroid tissue was incubated in Krebs-Ringer bicarbonate (KRB) buffer (98.5 mM NaCl, 4.9 mM KCl, 1.2 mM KH_2_PO_4_ 1.2 mM MgSO_4_- 7H_2_O, 20 mM HEPES, 2.6 mM CaCl_2_-2H_2_O, 25.9 mM NaHCO_3_, pH 7.4) supplemented with 5 mM U-^13^C-glucose. This buffer was equilibrated at 37°C, 21% O_2_, and 5% CO_2_ prior to incubations. We incubated freshly dissected tissue in media at 37°C, 21% O_2_, and 5% CO_2_ for amounts of time indicated in each figure. Following each incubation, tissue samples were flash frozen in liquid N_2_. Medium was sampled every 30 minutes to evaluate glucose and lactate content.

### Glucose and lactate assays

Glucose and lactate levels in cell culture supernatant were determined enzymatically, as in (Seidemann, 1973) but adapted for use in a 96 well plate. A detailed description of the adapted glucose assay protocol can be found at https://www.protocols.io/view/glucose-concentration-assay-hexokinase-g6pdh-metho-dm6gpj5jdgzp/v1. A detailed description of the adapted lactate assay can be found at https://www.protocols.io/view/lactate-concentration-assay-ldh-method-6qpvr4733gmk/v1. For each assay, levels of glucose or lactate are determined by measuring absorbance of light at 340 nm, which reflects NADH or NADPH produced in the assay. We measured absorption of 340 nm light using a Synergy 4 plate reader (BioTek).

### Metabolite extraction

Metabolites were separated from tissue in an extraction buffer consisting of 80% methanol, 20% H_2_O, and 10 μM methylsuccinate (Sigma, Cat#: M81209) as an internal standard. The extraction buffer was equilibrated on dry ice, and 150 μL was added to each sample. Tissues were then disrupted by sonication and incubated on dry ice for 45 minutes to precipitate protein. Proteins were pelleted at 17,000 × g for 30 minutes at 4°C. The supernatant containing metabolites was lyophilized at room temperature until dry and stored at −80°C until derivatization. The pellet containing protein was resuspended by sonication in RIPA buffer (150 mM NaCl, 1.0% Triton X-100, 0.5% sodium deoxycholate, 0.1% SDS, 50 mM Tris, pH 8.0) and protein was determined by a BCA assay (Thermo Fisher, Cat#: 23225).

### Metabolite derivatization for Gas Chromatography-Mass Spectrometry

Dried samples were derivatized with 10 μL of 20 mg/mL methoxyamine HCl (Sigma, Cat#: 226904) dissolved in pyridine (Sigma, Cat#: 270970) and incubated at 37°C for 90 minutes. Samples were further derivatized with 10 μL tert-butyldimethylsilyl-N-methyltrifluoroacetamide (Sigma, Cat#: 394882) and incubating at 70°C for 60 minutes.

### Gas chromatography-mass spectrometry

Metabolites were analyzed on an Agilent 7890/5975C GC-MS using methods described extensively in previous work (Chertov et al., 2011; Du et al., 2015). One microliter of derivatized sample is injected and delivered to an Agilent HP-5MS column by helium gas (flow rate: 1 mL/min). The column temperature starts at 100°C for 4 minutes then increases by 5°C/min to 300°C, where it is held for 5 min. We collect ion intensities after a 6.5-min solvent delay.

Selected ion monitoring (SIM) records only selected m/z in expected retention time windows. These masses range from m/z: ∼50–600. Peaks were integrated in MSD ChemStation (Agilent Technologies, version E.02.01.1177), and correction for natural isotope abundance was performed using IsoCor 1.0 (Millard et al., 2012; Heinrich et al., 2018). Corrected metabolite signals were converted to molar amounts by comparing metabolite peak abundances in samples with those in a ‘standard mix’ containing known quantities of metabolites. Multiple concentrations of this mix were extracted, derivatized, and run alongside samples in each experiment. These known metabolite concentrations were used to generate a standard curve that allowed for metabolite quantification. Paired tissues such as retinas and RPE-choroid from the same mouse were treated as technical replicates and averaged.

### Liquid chromatography-mass spectrometry

For liquid chromatography, samples were separated on an Acquity 2D UPLC equipped with an Acuity BEH Amide column (Waters, 2.1 × 50 mm, 1.7 μm). The mobile phases were (A) water with 10 m*M* ammonium acetate (pH 8.9) and (B) acetonitrile/water (95%/5%) with 10 m*M* ammonium acetate (pH 8.9). All solvents were LC–MS Optima grade (Fisher Scientific). The protocol was 0 min: [5% A, 95% B], 2 min [15% A, 85% B], 2.8 min: [40% A, 60% B], 3.2 min: [50% A, 50% B], 3.7 min: [15% A, 85% B], 4.4 min: [5% A, 95% B], at a flow rate of 0.4 mL/min. Chromatographically separated glucose was quantified on a Waters Xevo TQ mass spectrometer with electrospray ionization. Glucose ions were collected in negative mode with a cone = 18 V, collision energy = 8 V, parent ions of m/z = 179-185 and daughter ions of m/z=89-92. Dwell time for each ion was 0.08 s.

### Statistical Analysis

All displayed data include mean ± standard error and may additionally show individual experimental replicates. Statistical analyses were performed using Prism (GraphPad Software, version 10.0). Significance thresholds were set to α<0.05. When p-values are not directly shown, * indicates p<0.05, ** indicates p<0.01, *** indicates p<0.001. “n” signifies sample size, which is indicated in figure legends for corresponding experiments. For glucose depletion and lactate production in medium (**Fig. 2**), a linear regression was used to calculate slopes for glucose consumption or lactate production over the first hour of the experiment. Slopes were compared using a student’s t-test. T-tests were also used to analyze the data in **Fig. 4A-L**. Other data were analyzed by a two-way ANOVA (**Fig 3. C-E**, **Supplemental Fig. 3**). Post-hoc comparisons for 2-way ANOVAs and multiple T-tests (**Fig. 4**) were made using the two-stage step-up method of Benjamini, Krieger, and Yekutieli.

## Notes

### Competing Interest Statement

The authors have declared no competing interest.

### Summary of Updates

This has been substantially revised and edited for the sake of clarity and to improve the validity of the scientific conclusions.

