## Supplementary figures and images for "*In vivo* exchange of glucose and lactate between photoreceptors and the retinal pigment epithelium"

### Supplemental Fig 1

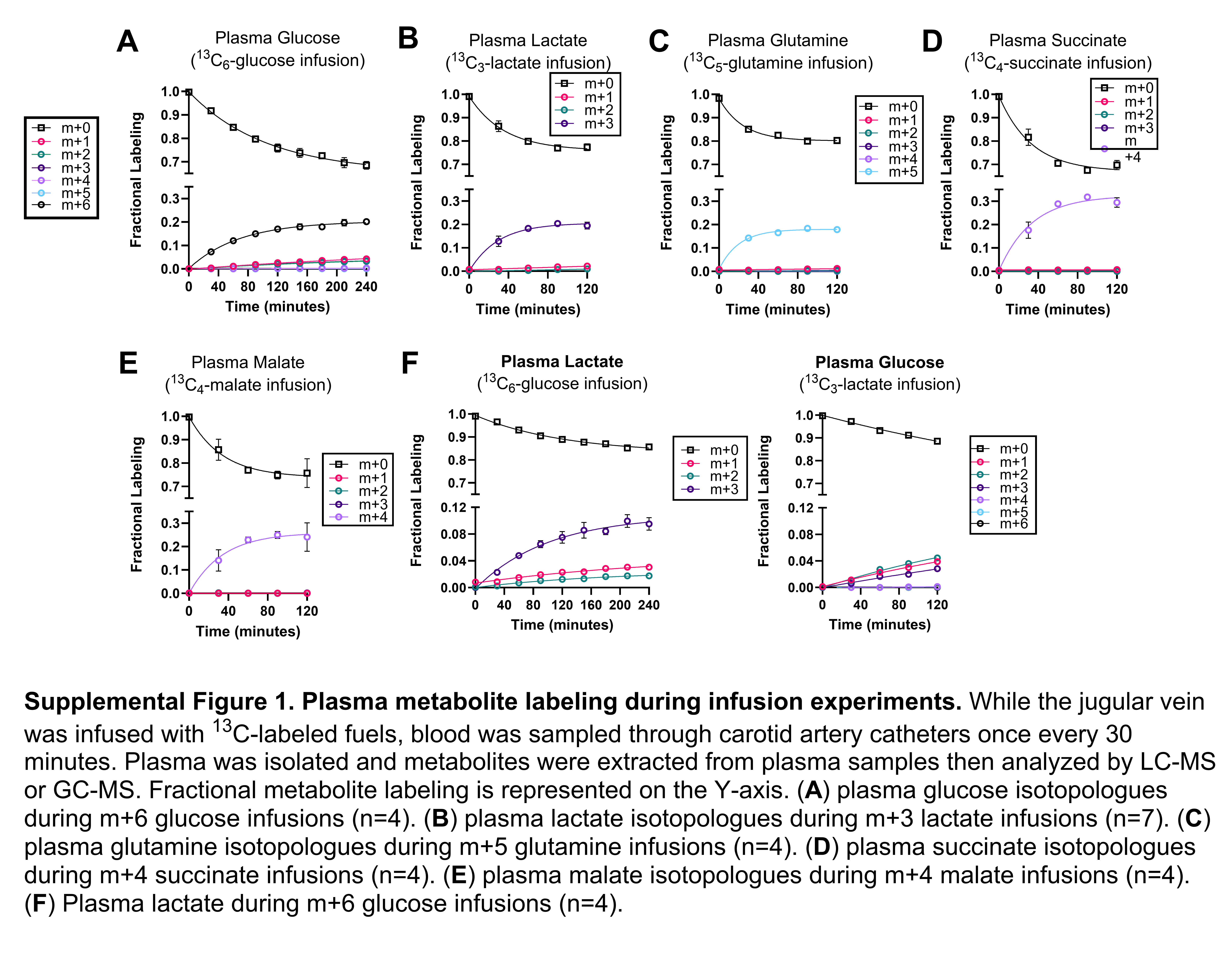

### Supplemental Fig 2

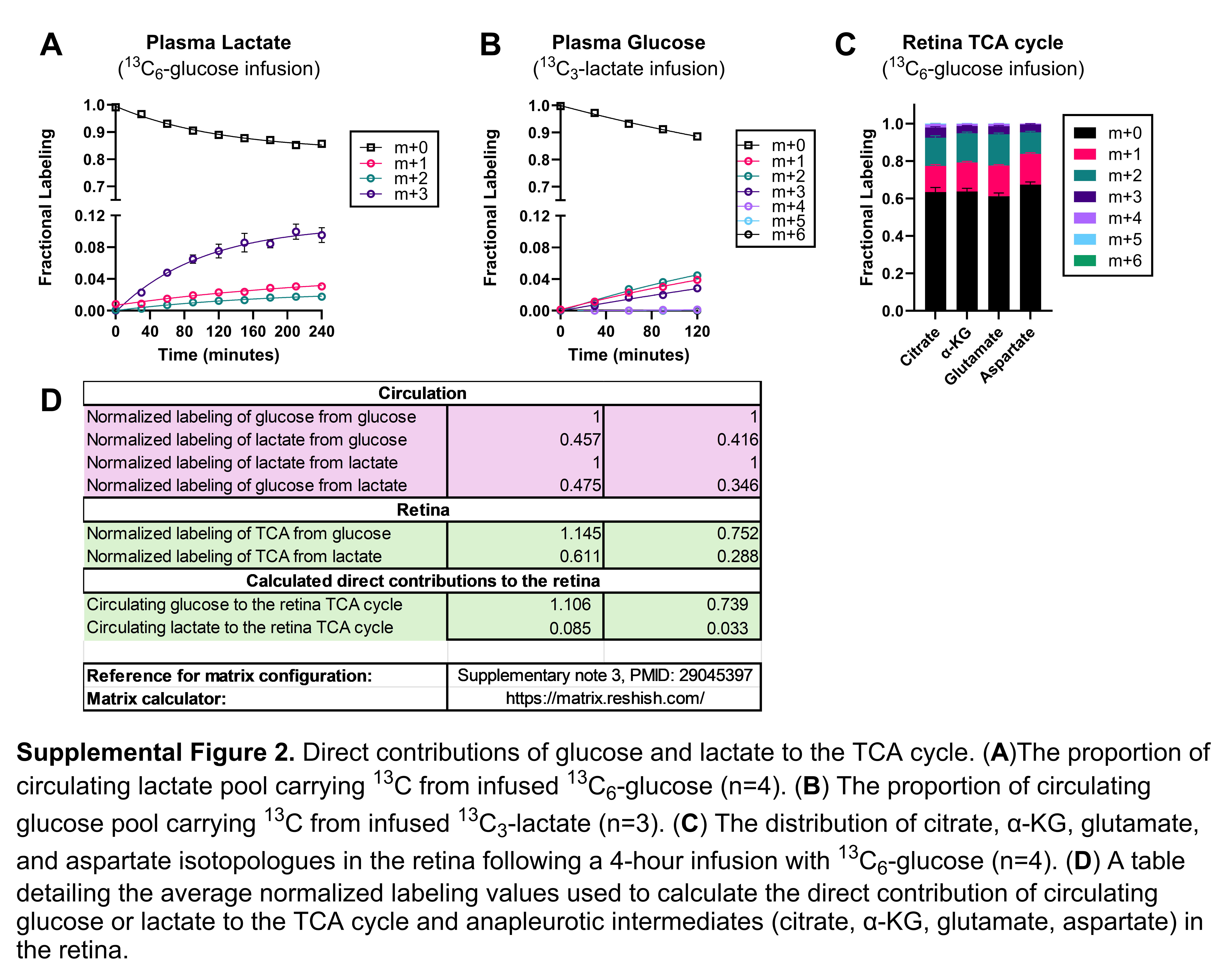

### Supplemental Fig 3

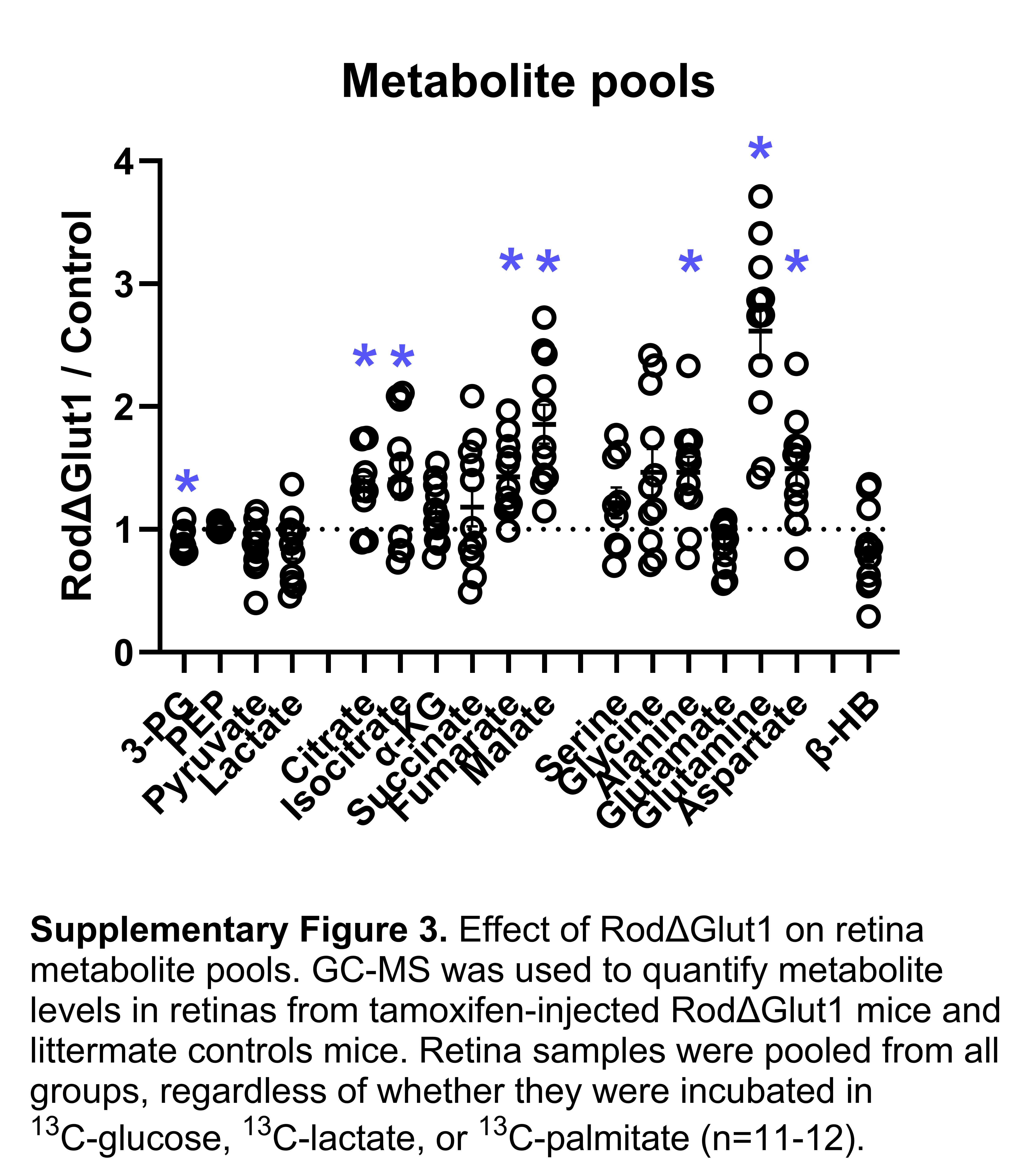

### Supplemental Fig 4

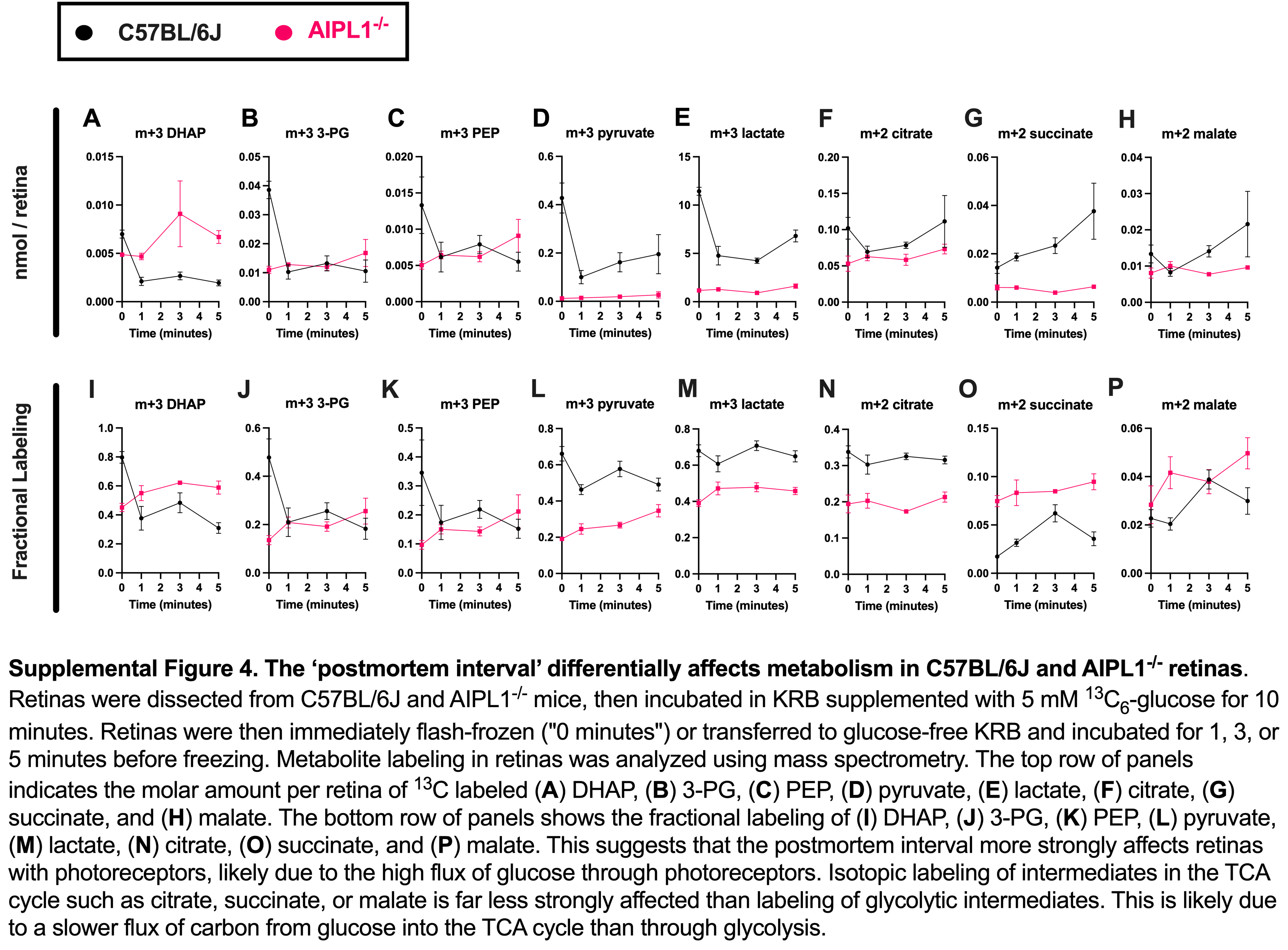
